# Brain Data Standards - A method for building data-driven cell-type ontologies

**DOI:** 10.1101/2021.10.10.463703

**Authors:** Shawn Zheng Kai Tan, Huseyin Kir, Brian D. Aevermann, Tom Gillespie, Nomi Harris, Michael Hawrylycz, Nik Jorstad, Ed Lein, Nicolas Matentzoglu, Jeremy A. Miller, Tyler S. Mollenkopf, Christopher J. Mungall, Patrick L. Ray, Raymond E. A. Sanchez, Brian Staats, Jim Vermillion, Ambika Yadav, Yun Zhang, Richard H. Scheuermann, David Osumi-Sutherland

## Abstract

Large-scale single-cell ‘omics profiling is revolutionising our understanding of cell types in complex organs like the brain, where it is being used to define a complete catalogue of cell types, something that traditional methods struggle with due to the diversity and complexity of the brain. But this poses a problem. How do we organise such a catalogue - providing a standard way to refer to the cell types discovered, linking their classification and properties to supporting data? Cell ontologies provide a solution to recording definitions, classifications, and properties of cell types and provide standard identifiers for annotation, but they currently do not support the data driven cell type definitions and classifications needed for multi-modal single cell ‘omics profiling.

Here we describe the construction and application of a semi-automated, data-linked extension to the Cell Ontology that represents cell types in the Primary Motor Cortex of humans, mice and marmosets. The methods and resulting ontology are designed to be scalable and applicable to similar whole brain atlases currently in preparation.

## Introduction

The large-scale application of omics profiling techniques at the single-cell level is producing enormous volumes of data. Cell ontologies are poised to play a critical role in making these data searchable and integratable ^1^. At the same time, the application of these profiling techniques is revolutionising our understanding of cell types and cellular heterogeneity ^2,3^. The impact of this revolution is especially dramatic for the brain. Due to the complex cellular architecture of the brain, traditional qualitative, categorical methods of classifying neurons based on location, morphology, marker expression and function have not achieved a coherent, unified view of granular brain cell types and their classifications. This has begun to change with the application of massively parallel single-cell or nucleus RNA sequencing (sc/snRNAseq) methods to the brain, combined with multimodal transcriptomic techniques such as Patch-seq ^4^. The BRAIN Initiative Cell Census Network (BICCN) recently completed a comprehensive, multimodal cell census and atlas of the primary motor cortex across multiple species ^5–7^. This takes the approach of treating consensus clustering of similar cells from single nucleus RNA-seq data from multiple experiments as a ground truth for defining cell types and their classification.. The resulting cell type hierarchies serve as anchors for alignment of data from other modalities, allowing spatial localization, morphology, electrical properties, chromatin accessibility, and other features of cell types to be recorded and compared across species. Evidence from systems in which a more comprehensive classification of cell types has been achieved by classical methods than has been possible in the brain suggests that the classifications resulting from sc/snRNAseq analysis align closely with classically defined types ^8^.

This poses challenges for standard approaches to ontology development. How are we to integrate cell types defined with reference to clusters of transcriptomically similar cells into cell ontologies in which cell type/classes are defined using simple, categorical assertions about their morphological and functional properties, location and marker expression? How can we do this in a way that is transparent about the origins and evidence for these classifications? How can we enable ontology users to leverage the data used to define and classify reference cell types in the ontology to classify cell types represented in their own data?

Here we describe a solution to these challenges in the form of a template-driven ontology generation pipeline and an ontology of cell types defined in the BICCN mini-atlas, Brain Data Standards Ontology (BDSO), that forms part of the Provisional Cell Ontology ^3^, which extends the Cell Ontology ^9^ with potential new cell types from single cell analysis. Ontologies should serve as both an easily searchable source of terms for annotation and a data structure supporting organisation, search and navigation of annotated data. We demonstrate the utility of our ontology for this via its application to the organisation, search and navigation of data about cells in the mini-atlas on the Allen Cell Type Explorer web app.

## Results

### Brain Data Standards Ontology Design

One of the outputs of the BICCN mini-atlas ^10^ is a standardized representation of cell clusters (CCN) and the hierarchical relationships between them that constitute the ground-truth for cell-types defined in the atlas. The clusters and their hierarchical arrangement derive from unsupervised, hierarchical clusterings of single-cell transcriptomic and epigenetic profiles of the primary motor cortex in mouse, human, and marmoset ^10,11^. Each individual hierarchical clustering (referred to here as a taxonomy) is either created from a single data set (e.g., in marmoset) or through a consensus of two (human) or many (mouse) data sets. Using mouse transcriptomics clusterings as an anchor, morphological and electrophysiological profiles of single-cells are mapped to omics-based types using Patch-seq data ^7^. Finally, comparison of clusters across species is used to generate cross-species mappings and groupings of clusters which represent putative homology groupings ^10,11^. All of this information is available in a standard format (common cell type nomenclature taxonomy files, here referred to as CCN taxonomy files) developed by the BICCN to represent mammalian brain cell type taxonomies and the relationships between them ^12^.

To produce a set of definitional characteristics of the cell types identified in these taxonomies, a minimum set of markers that can be used to distinguish cells in that cluster from those in other clusters in the same taxonomy was produced using the NS-Forest algorithm ^13^. Taking the clusters as ground truth for all cell types present in the primary motor cortex, the combined expression of each marker set should be necessary and sufficient to identify the corresponding cell type in the context of the primary motor cortex.

The BDSO is built as a faithful representation of the BICCN mini-atlas cell type taxonomies (Figure 1). In order to achieve this, we first devised a schema to represent taxonomies in Web Ontology Language, OWL2 ^14^, the formal language we use for constructing ontologies. OWL2 makes a distinction between individuals, e.g., an individual neuron depicted in a micrograph, and classes, e.g., the class of all Chandelier neurons. Each taxonomy is represented in BDSO as a collection of OWL Individuals, with each Individual representing a cluster of single-cell transcriptomes and retaining all original metadata in the CCN taxonomy file from which it is derived. Hierarchical clustering is represented by relating these individuals to each other via a transitive subcluster_of relation.

**Figure 1.**
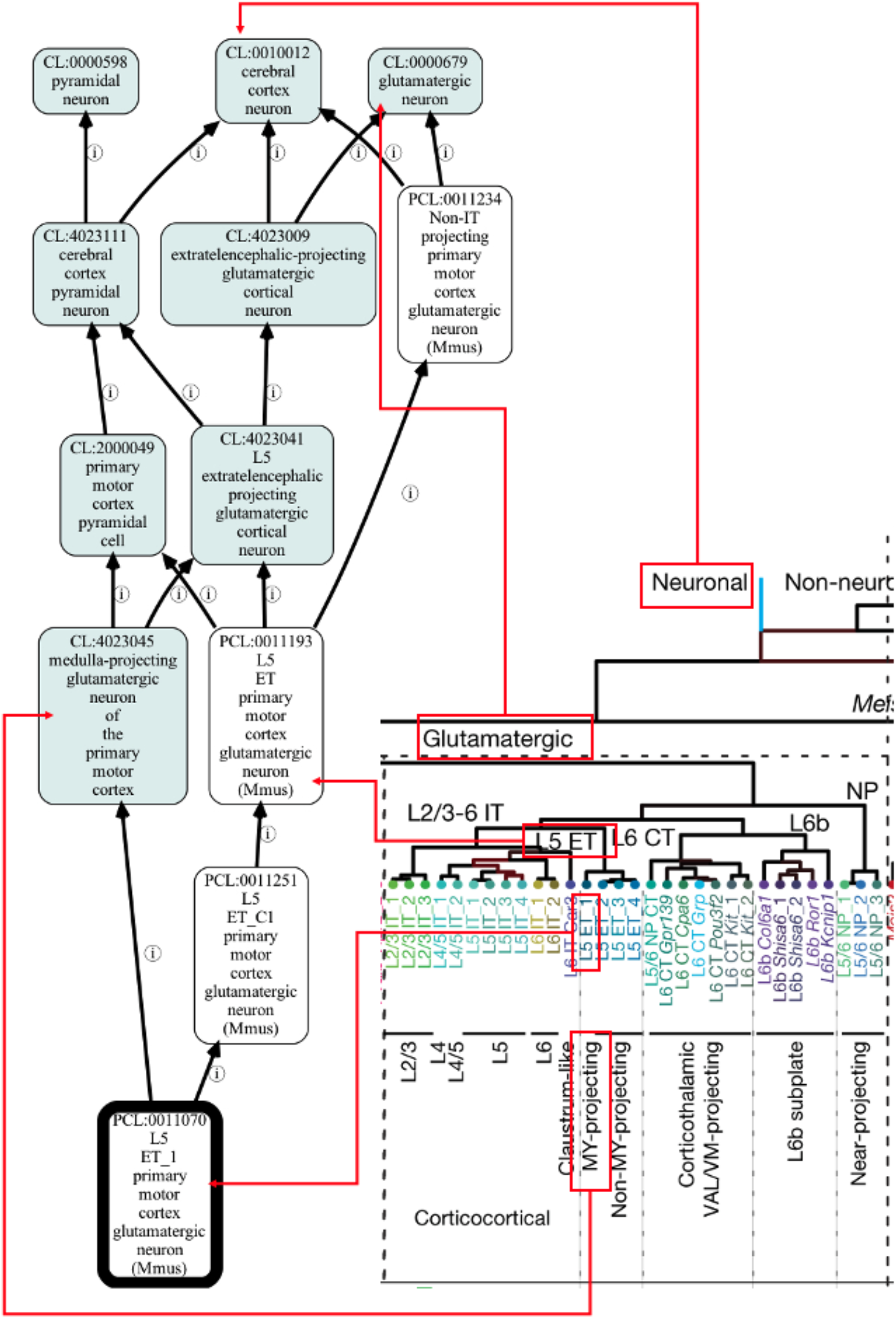
Example of representing the BICCN mini-atlas cell type taxonomy in an ontology. Red boxes/lines show how terms in the taxonomy are mapped into an ontology format (visualised by the Ontology Access Kit).

Each taxonomy has many more nodes than it would be reasonable to create classes for. In order to select useful intermediate nodes for representation, taxonomy authors of the BICCN mini-atlas flagged nodes to generate a 3-level hierarchy with the most granular level consisting of all leaf nodes ^10^. We generated cell classes for all tagged clusters, apart from some high-level groupings (e.g. all cells, non-neuronal, etc.) that would not make sense as a cell type term as they are overly generalised. Each of these classes is linked formally to a cluster individual using a standard pattern in OWL that can be used by standard OWL reasoning software to automatically build a classification hierarchy for the BDSO classes (see Fig. 2 and the next section for more details). Lastly, we treated cross-species mappings between cell types as putative homology mappings, by using the relation in_historical_homology_relationship_with ^15^ (imported from the OBO relations ontology) in a pairwise manner.

**Figure 2.**
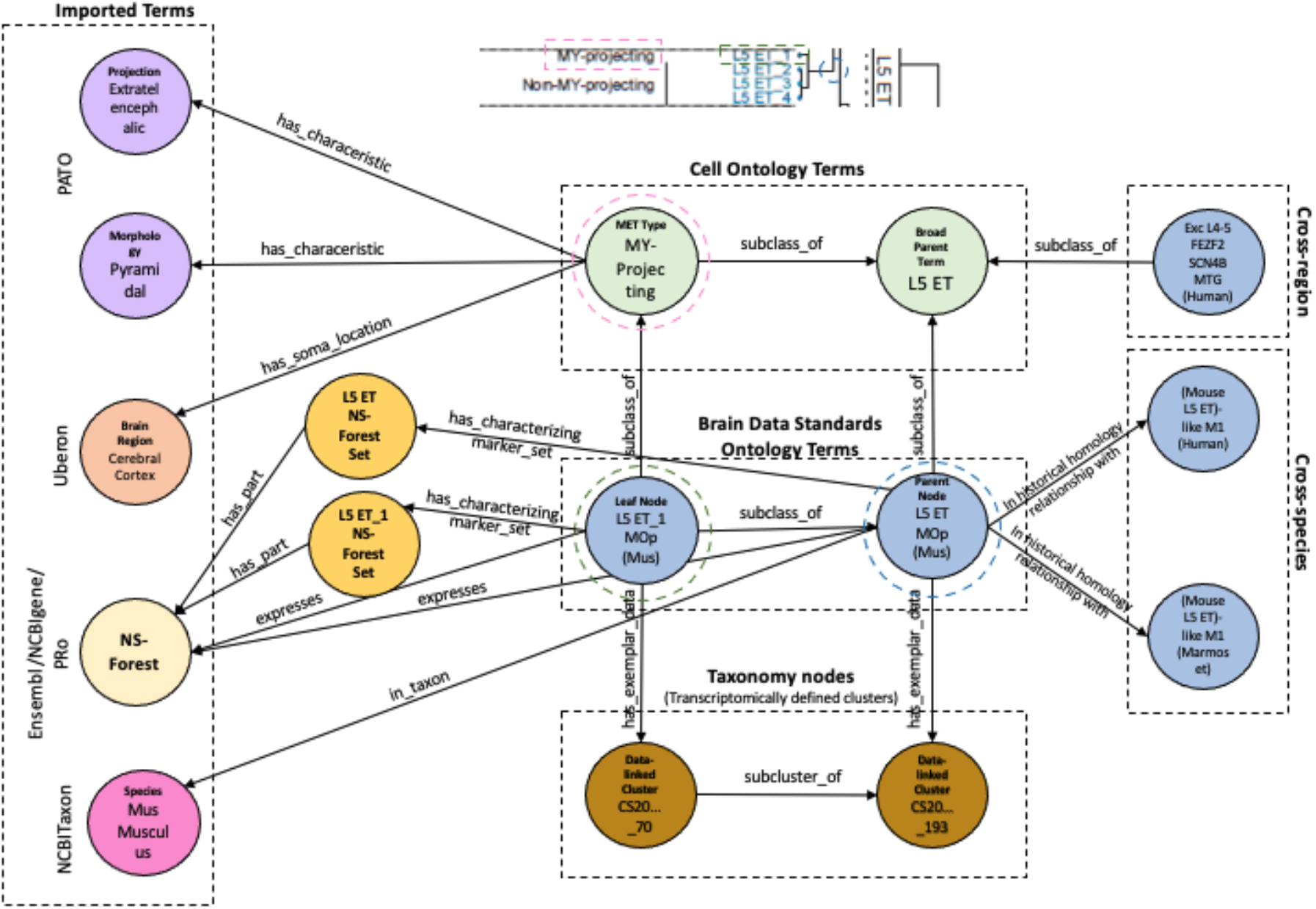
Graph illustrating the BDSO schema. This graph shows the relationship of the BDSO classes (Brain Data Standards Ontology nodes, light blue circles) to OWL Individuals (Taxonomy nodes, brown circles) representing clusters in the data-driven taxonomy used as input and to the build process, to classes in the Cell Ontology (green circles) and from external ontologies (imported terms box) representing species (NCBITaxon), brain region (UBERON), morphology (PATO), and markers (Ensembl/PRO). NS-Forest marker combinations are represented through sets, with individual markers being part_of them. The right side of the figure shows links to potentially homologous cell type classes (Cross-species box) using the relation (OWL objectProperty) ‘in historical homology relationship with’ and cross-region terms (Cross-region box).

To integrate the BDSO with existing ontologies, classes defined for intermediate nodes in the hierarchy are further classified using classes in CL, which we have extended as required (e.g., see ‘L5 extratelencephalic’ class in Figure 2). These include classes that are defined by expression of classical marker genes (e.g., VIP-expressing GABAergic neurons), morphology (pyramidal) or projection pattern (extratelencephalic projecting), mapped based on co-collected transcriptomic profiles ^10^. The BDSO also reuses existing ontologies to represent species (NCBITaxon ^16^), brain region (UBERON ^17^), morphology (PATO), and marker genes (Ensembl/PRO ^18,19^). All relationships added use OBO standard relations from the OBO relations ontology and follow or extend standard schemas used by CL (Figure 2). In addition to tightly integrating these terms with CL, this approach maximises the potential for making data annotated with BDSO interoperable with the many other datasets annotated with these ontologies.

### Designing an automated pipeline

Manually building an ontology to represent the huge amount of data from the BICCN mini-atlas is impractical, error-prone, and unscalable. It was therefore imperative to harness automated tools to build the BDSO. To build the BDSO, we use CCN taxonomy files, NS-Forest marker gene mappings and reference gene lists as input to a semi-automated pipeline. The pipeline takes advantage of the schema described in Figure 3 to build a hierarchy that mirrors the cluster hierarchy (see L5 ET in Figures 1 and 3 for example implementation). The BDSO is built using the Ontology Development Kit ^20^ and uses standard ontology term templating systems ^21,22^ to generate labels, definitions and synonyms for BDSO terms and to add CL classifications and relationships^1^ recording location (using Uberon terms ^17^), species (using NCBI taxonomy terms ^16^), markers, projection patterns and morphologies (see Figure 4 for examples). The results of NS-Forest analysis, ingested via standardised TSV files, are automatically consumed by the pipeline and integrated into the ontology (see section below). Manual curation such as mapping to CL terms, adding cell properties (morphology, projections, etc.) were kept to a minimum and done via templates to ensure consistency and scalability.

**Figure 3.**
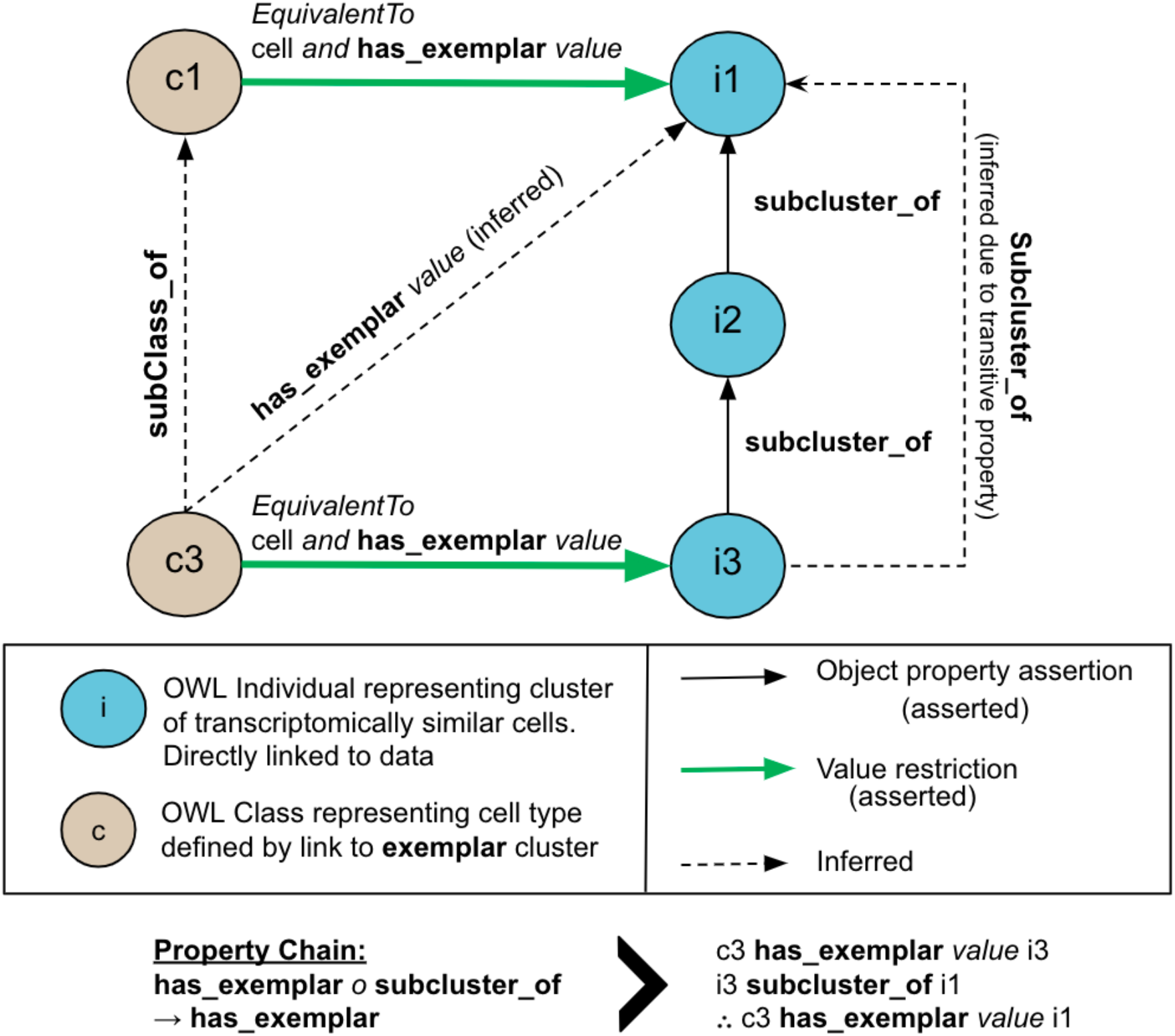
Representative schema for data-driven classification. Blue nodes (i1-3) are OWL individuals representing clusters of single-cell transcriptomes, while tan nodes (c1, c2) are OWL classes representing cell types. Hierarchical clustering is represented using the transitive subcluster_of relation (objectProperty) to link individuals. Each class is defined by reference to a cluster individual (i), via the relation (objectProperty) as equivalent to (any) cell that has_examplar (value) i. Reasoning via a chain of these two properties (bottom and right sides of the diagram above) is sufficient to infer that c3 has_examplar value i1 and so, combined with the assertion that it is a (type of) cell, fulfils the conditions required to be a subclass of i1.

**Figure 4.**
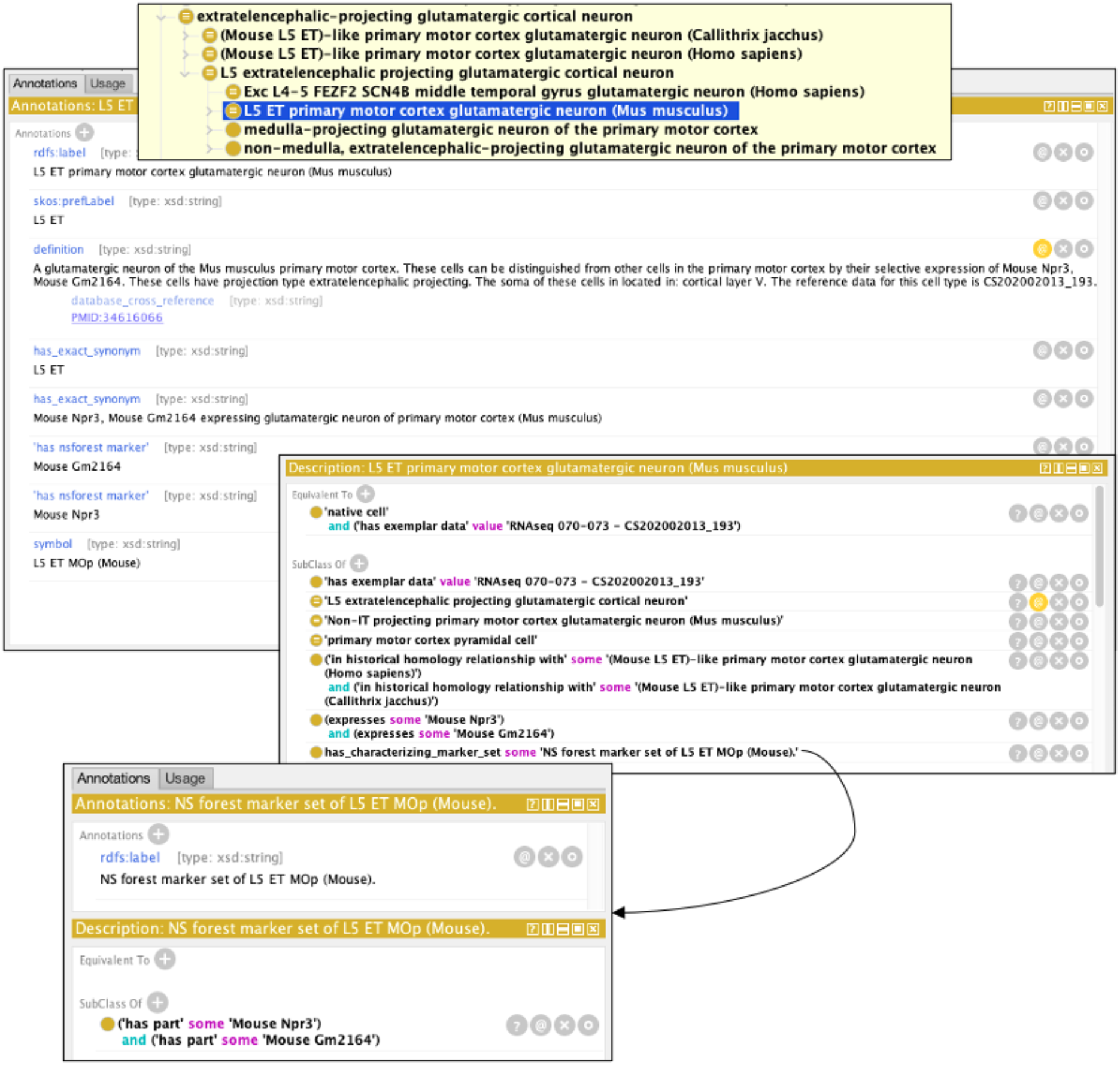
Example of an automatically generated class displayed in the Protege ontology browser. In this example, we show L5 Extratelencephalic (ET), which is a grouping class. The label, definition, and set of synonyms are auto-generated from OWL templates using a Dead Simple OWL Design Patterns (DOSDP) system. Automatic axiomatisation includes brain region, species, NS-Forest markers, projection pattern, morphology, named markers, and has_exemplar_data link to taxonomy node (cluster), using a reification pattern. This results in the reasoner classifying this class under L5 extratelencephalic projecting glutamatergic cortical neuron (based on automated axiomatisation of brain region and projection pattern), and primary motor cortex pyramidal cell (based on automated axiomatisation of morphology and brain region). has_characterzing_marker_set schema for NS-Forest is also shown.

### Representing data and analysis results

The BDSO uses the direct results of data analyses as evidence for the existence of cell type classes. To reflect this, and to allow users direct access to the data that justifies the categorical assertions that we make, we link the ontology clusters to datasets (expression matrices) available on Nemo (https://assets.nemoarchive.org/dat-ch1nqb7), and we include the quantitative data that support categorical assertions made in the ontology, where this data is available. Currently, we include a measure of the accuracy of classification using NS-Forest marker F-Beta scores and we plan to incorporate measures of transcriptomic similarity to support homology assertions. CCN taxonomy files include a measure of confidence in the division into (sibling) subclusters, plotted as height in dendrogram views. We retain this measure, along with all other metadata, attached to individual clusters.

Each set of NS-Forest markers should theoretically be necessary and sufficient for identifying a cell type with high precision within the dataset used to define them. In the case of the mini-atlas, the datasets correspond to all cells with a soma located in the primary motor cortex of some specified species and so should be necessary and sufficient for identifying the cell type within that anatomical context more generally. We also have evidence that they are useful for detecting the same cell type in other brain regions: In many cases, the markers identified by NS-Forest in the primary motor cortex, are expressed in equivalent cell types found in another cortical brain region (middle temporal gyrus) ^23^ however the NS-Forest algorithm typically finds other sets of makers in these cases.

We record this context as a restriction on the class using a has_soma_location to the brain region and represent NS-Forest markers through an NS-Forest set class, ‘S’ in the example below, with marker genes as parts (See Figures 1 and 3):

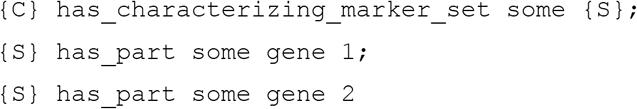

This approach allows us to record multiple marker sets for each cell type, which may be essential in future, given the many competing methods available for defining cell type markers. The intermediate node allows for clear grouping of marker sets in knowledge graphs (see Figure 2). We also use the node to record Fβ scores for each set - recording the accuracy of classification using the markers on the reference transcriptomic datasets. We do this through a custom annotation property ‘fbeta_confidence_score’ that is annotated on the marker set class.

We rejected an alternative approach, of using an EquivalentClass axiom with clauses to restrict for location and NS-Forest markers to formally specify necessary and sufficient conditions, as having two equivalence axioms to define a cell type can potentially lead to competing classifications.

### Ontology content summary

The latest release (2022-04-27 Release) of the BDSO component (which PCL imports) contains 913 individuals, out of which 890 are taxonomy nodes (individuals also include datasets), and 112447 classes (including genes and NS-Forest sets), out of which 1384 have the PCL namespace and 555 are cell types. The remaining terms are imported from OBO ontologies into PCL. All object properties used are imported from RO as per OBO foundry guidelines.

### Application

A key function of the BDSO is to support organisation, navigation and searching of data in a community-accessible view of the cell types defined in the BICCN mini-atlas of the mammalian primary motor cortex ^10^ through a web-based application (web-app) that integrates cell type descriptions and related data, known as the “Cell Type Knowledge Explorer” (Figure 5). Each page in this web-app corresponds to a cell type defined with reference to a cluster in one of the BICCN taxonomies represented in the BDSO, and features a wide range of data and analysis from multiple cross integrated datasets. The aim of the ontology-driven search and navigation tools is to support access to these pages in the web-app.

**Figure 5.**
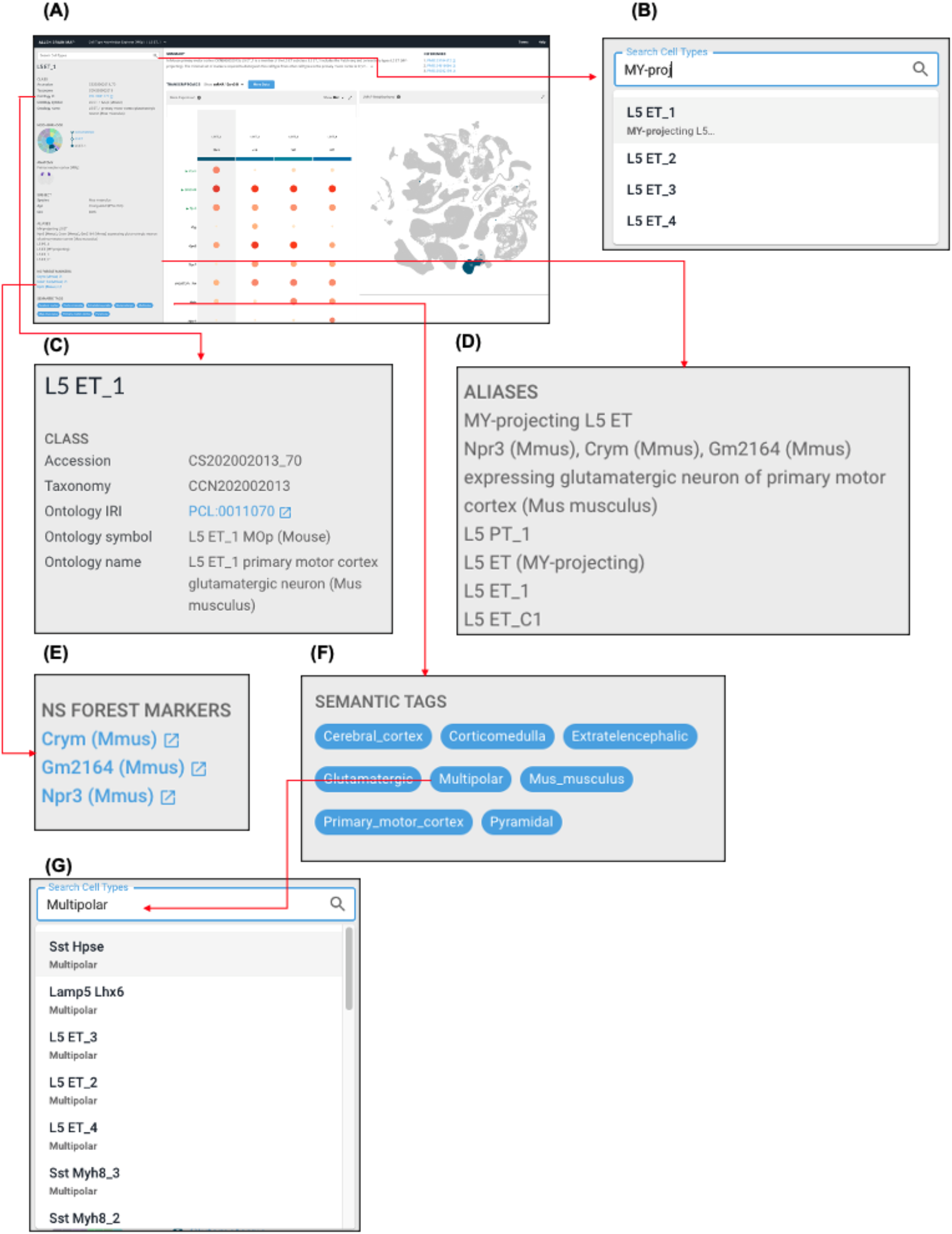
Screenshots of the alpha version of the Cell Type Knowledge Explorer web app, incorporating search and navigation functionality driven by the BDSO. (A) An overview of the web app with the ontology incorporated into it. Red arrows show zoomed in version and directional links. (B) An example of autocomplete search, which also allows search by synonyms. (C) Information about the cell type incorporates ontology identifiers, ontology symbols, and ontology names. (D) A list of synonyms generated by ontology annotations and extra curated synonyms. (E) A list of NS-Forest markers with links out to their identifiers.org pages. (F) Semantic tags of the cell type corresponding to species, brain region, and cell properties such as morphology (pyramidal) and projection pattern (extratelencephalic). Clicking on one of these panels drives faceted search through the search bar seen in (G).

While expressiveness of ontology formats such as OWL is an advantage for semantic data processing, OWL is complicated to develop applications with and has limited tooling. Graph databases like neo4j, and indexed document stores such as SOLR and ElasticSearch, provide a more tractable, fast way to drive web applications. For this purpose, we extended a library, neo4j2owl ^24^, developed for the Virtual Fly Brain project ^25,26^, that ensures logical projection of OWL ontologies into labelled property graphs. Neo4j2owl imports OWL ontologies into Neo4j in a way that preserves entailments and annotations, but not the syntactic complexities of OWL. It also supports the addition of semantic tags, in the form of simple strings attached to classes and individuals, driven by OWL DL or SPARQL queries. We use this semantic tag system to provide an application-specific, gross classification that provides additional information about classes in a useful form to users and can be used to drive faceted search. For example, we can tag all classes corresponding to subclasses of GABAergic neuron, or all classes fulfilling an OWL DL query for classes of neuron with pyramidal morphology (see Figure 5f). The full Knowledge Graph can be accessed at http://purl.obolibrary.org/obo/pcl/bds/kg/.

An illustration of the resulting property graph is shown in Figure 2. These property graphs allow applications such as the cell type knowledge explorer to use the ontology data to populate parts of the application and enable full-text and faceted search functions.

Ontology-based navigation and search functions are provided through two mechanisms - autocomplete (which takes advantage of curation of synonyms in the ontology) and faceted search (Figure 5). Autocomplete allows users to search for cell-type ontology terms, displaying a list of lexical matches for users to choose from (Figure 5b). Faceted search of Cell Type Knowledge Explorer works via a set of tags corresponding to gross classifications (e.g. GABAergic), intrinsic properties (e.g. pyramidal morphology) and extrinsic properties (brain region location, species) of cell types, added to cell type neo4j nodes via OWL DL queries of the underlying ontologies. Currently, implementation of this works through automatically adding the term to the search bar and allowing the free-text search to complete the search (Figure 5F,G). However, this approach is unlikely to scale as the content of cell type explorer grows. There are plans to allow users to take better advantage of faceted browsing using semantic tags via a results page that can be refined via combinations of semantic tags combined with lexical search, allowing users to find neurons by any combination of location, morphology, species, neurotransmitter and name/synonym substring.

## Discussion

The BDSO is a faithful representation of the data-driven, consensus cell type classification that includes the BICCN mini-atlas of the mammalian motor cortex ^10^. By using a schema that defines classes logically via links to an OWL representation of data and analyses, we can use OWL to directly leverage the data-driven taxonomy of the miniatlas to classify cell types in BDSO using OWL reasoning. As a result, classes retain direct links to the data and analyses that define them and the origins of this classification are transparent and insulated from the manual editing process that might alter or obfuscate them. Using templated specification of ontology classes, the BDSO build process is scalable and extensible and allows a flexible mix of automation and manual curation. It also makes it possible to update as new, improved versions of data-driven classifications of the same cell types are released. The linked data can potentially be used to replicate analyses and to map cell types defined in BDSO to other datasets (e.g., using Azimuth ^27^, FR-match ^23^). The addition of NS-Forest markers ^13^, representing minimal markers for distinguishing, with high confidence, cell types from other cell types defined in the analysis, provides a simple mechanism for mapping cell types from third-party transcriptomics data to the BDSO.

In future, we plan to incorporate measures of transcriptomic similarity in support of homology assertions and a measure of confidence for data-driven taxonomy nodes. We will also incorporate contextual information about the nature of these measures. While the absolute values of these measures are inevitably specific to the datasets/analysis they come from, they are at least usable for intra-dataset comparisons.

As a broader consensus and whole-brain datasets emerge, we expect NS-Forest F-Beta scores and taxonomy node confidence measures to be informative of which cell types we consider stable and replicable.

While the approach described meets many of the requirements for a scalable approach to cell type representation, some challenges remain. The current representation lacks links to transcriptomic data from Patch-seq data used to map morphologically defined types. Using transcriptomic clustering as ground truth for an ontology also comes with its inherent challenges. Penetrance of marker expression and location to a specific cortical layer varies across clusters, so all/some quantified assertions of marker expression in OWL will always be an approximation and will always require either automated or qualitative assessment of thresholds. Finally, nomenclature issues frequently arise when data-driven classifications are mapped onto classically-defined classes. For example, the literature is full of references to VIP-expressing GABAergic neurons, identified using VIP as a marker, but clustering defines a broader group of related GABAergic neurons including some subtypes that do not express VIP, at least not at levels detectable by snRNAseq in the adult.

The transcriptomic approach potentially allows the definition of transcriptomically defined, species-neutral grouping classes. We decided against adding these because the resulting classifications are not likely to remain stable as more species are added to the analysis, although this may change in future with large-scale analyses using many species. It is also likely to be challenging to map these classes to the more traditionally defined species-neutral cell type ontology classes.

Another challenge comes from working with nomenclature defined by researchers. Terminology that makes sense in the limited local context of a dataset can be confusing to users viewing it in the broader, integrated context of an ontology. In the primary motor cortex mini-atlas datasets used for this work, names given to cell types in human and marmoset were derived from the names of the mouse cell types, even where that name implies properties (e.g., marker expressions) that do not apply. For example, the Sncg cluster in marmoset is aligned to that of mouse Scng cluster but contains many cell types that do not necessarily express Sncg (Figure 6). To make this clear we rename these terms following the pattern mouse {x} like, e.g., (Mouse Sncg)-like (Marmoset).

**Figure 6.**
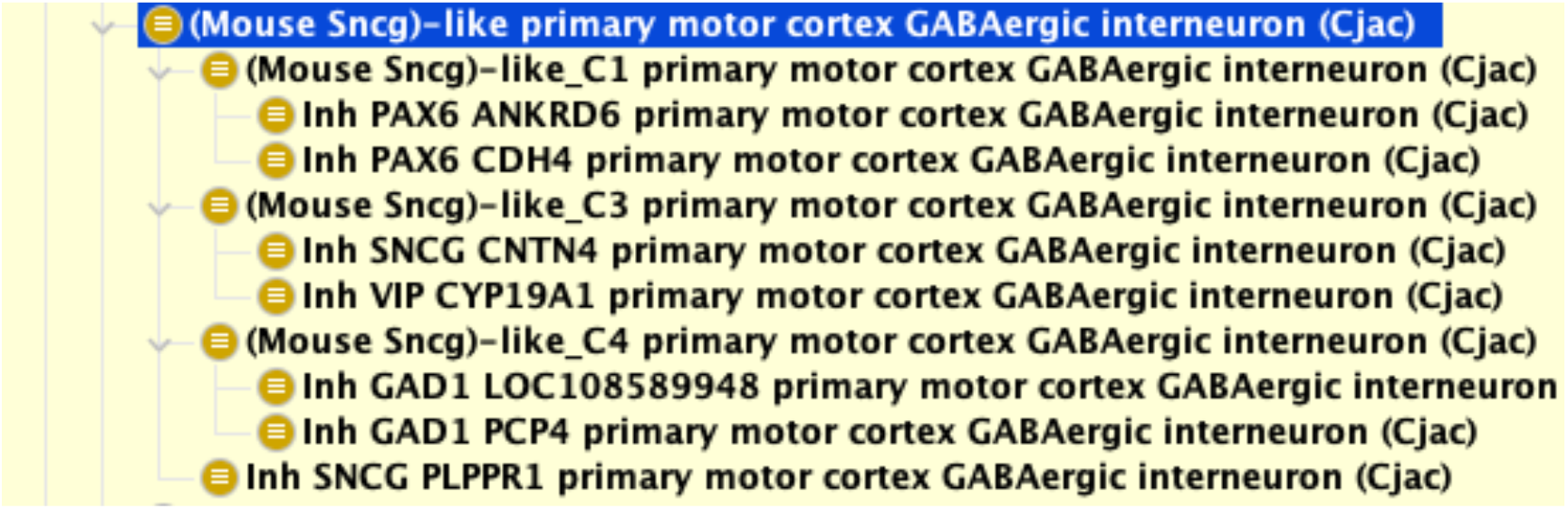
Example of a cell type name that is derived from the names of the mouse cell types. The Marmoset (Callithrix jacchus) cell type taxonomy is aligned to the mouse cell type taxonomy, resulting in a “sncg grouping” that contains cell types that do not necessarily express Sncg. To make this clear, the class was renamed (Mouse Sncg)-like.

Lastly, as efforts to expand scRNAseq cell typing to the entire brain, there is a crucial need for upstream standardisation and validation in order to efficiently scale up what we have presented in this paper. Tooling that allows biologists to annotate cell types with existing terms created through the BDSO, automated checks for quality control, and consensus on data formats, nomenclatures, and version control are all required if we are to effectively manage the huge input of data that is inevitable from such work.

## Conclusion

The BDSO acts as a functional tool for managing data from the BICCN mini atlas project, underlying the search and navigation of the Cell Type Knowledge Explorer web application, and provides a controlled vocabulary for future annotations. Beyond its practical function, it is also an example of how ontologies can harness automation to process the large amounts of data that is inevitable with the rise of sc/snRNAseq methods. Crucially, the work on the BDSO has highlighted the need for good tooling and integration into the early steps of the processes of sc/snRNAseq experiments. While questions on representation of confidence, nomenclature, and links to data still require addressing by both the ontology and neuroscience communities, the BDSO is a practical first step to ontologising taxonomies generated by sc/snRNAseq cell typing in the brain.

## Methods

### Data Source

Input to the ontology was derived from data from the BICCN mini-atlas ^10^ and scRNAseq of the human middle temporal gyrus ^28^. NS-Forest analysis was done as previously described ^13^ using gene lists available from either NCBI gene ^16^ or Ensembl ^18^.

### Development Strategy

BDSO is developed based on the OBO Foundry ^29,30^ and FAIR ^31^ principles. Ontology terms were reused as much as possible (see results section) with all relationships used coming from the relations ontology and design patterns following or extending those used in the Cell Ontology. The BDSO is fully compliant with OBO Foundry standards and has been included as an ontology in the OBO Foundry.

### Templating Systems

The templating systems used in the automated pipeline are ROBOT ^21^ (used to generate individuals) and DOSDP ^22^. Briefly, information is extracted from the CCN taxonomy files and translated into template files that are processed either through ROBOT templates to generate individuals, or template files for classes where a curator manually curates additional information (e.g. mappings to CL cell types, morphology, etc.) which is then processed, together with NS-Forest markers, using DOSDP. These files are then merged as part of the pipeline for the final product.

### Provisional Cell Onotlogy

We updated the Provisional Cell Ontology to follow OBO Foundry standards by using a pipeline based on the ontology development kit ^20^. Earlier, manually generated releases of PCL shared terms with the version described here, but used non-standard IDs and schema. In order to support mapping of data previously annotated with PCL and references to PCL terms in previous publications ^3,11,28^, we mapped all original IDs to current OBO standard persistent URLs, using OBO standard mappings for obsoleted terms..

### Endpoints

As well as being available for downloading from a persistent URL (http://purl.obolibrary.org/obo/pcl.owl) and available for browsing on widely used ontology platforms including the Ontology Lookup service and Ontobee, the BDSO can be searched and queried via a REST API (http://purl.obolibrary.org/obo/pcl/bds/api/). These endpoints encapsulate the representational complexities of the underlying knowledge and property graphs and serve the ontology in web-friendly formats such as JSON. Using these endpoints, users can search for ontology terms, access their details and navigate through the ontology using relationships between concepts. Solr is used at the backend to provide enhanced full-text search and reduced service response times. The created Solr indexes are published publicly (https://github.com/obophenotype/brain_data_standards_queries).

### BDSO analysis

Statistics of metadata of BDSO were done using SPARQL queries with ROBOT ^21^ on the BDSO component. SPARQL queries used can be found in the repository (https://github.com/obophenotype/brain_data_standards_ontologies/tree/master/src/sparql).

### Figures Generation

Figure 1 ontology visualisation was generated by using the Ontology Access Kit ^32^ and dendrogram section was provided by the BICCN ^10^. Figures 4 & 6 uses screenshots from Protege ^33^. Figure 5 uses screenshots from the Cell Type Knowledge Explorer web app (https://knowledge.brain-map.org/celltypes/).

## Code Availability

The BDSO is generated using a dedicated ontology build pipeline, built as an extension to the Ontology Development Kit ^20^, but released as a component of the PCL, with all terms having PCL IDs. Previous releases of PCL ^3,11,28^ represented some of the same cell types as the current release but used a different, less formal schema and a different ID system ^3,11,28^. We have obsoleted these terms and provided a mapping, within PCL,to replacement terms allowing continued support for previous work annotated using PCL terms.

The BDSO’s code base is available at GitHub (https://github.com/obophenotype/brain_data_standards_ontologies) including documentation of the full technology stack and details of the approach. The latest release of the ontology is available for download from http://purl.obolibrary.org/obo/pcl/bds/bds.owl and is hosted on the EMBL-EBI ontology lookup service (OLS) ^34^ at https://www.ebi.ac.uk/ols/ontologies/pcl. OLS provides ontology search, browsing, visualisation capabilities and enables web services driven programmatic access to the BDSO.

## Acknowledgements

This work was funded by NIMH:1RF1MH123220-01 - “A Community Framework for Data-driven Brain Transcriptomic Cell Type Definition, Ontology, and Nomenclature.” We thank Maryann Martone and Carol Thompson for their invaluable contributions to discussions of the work described here.

## Data Availability Statement

All data used in the BDSO is publicly available and can be found in the original papers as well as nemo archive links available in the ontology. Code and source data used to generate the ontology is publicly available at GitHub (https://github.com/obophenotype/brain_data_standards_ontologies and https://github.com/JCVenterInstitute/NSForest). Primary motor cortex taxonomies are also publicly available (https://github.com/AllenInstitute/MOp_taxonomies_ontology).

## Competing interests statement

All authors declare no competing financial interests or potential conflicts of interest.

## Footnotes

1 More strictly, existential restrictions in Web Ontology Language (OWL).

## Notes

### Competing Interest Statement

The authors have declared no competing interest.

### Summary of Updates

Manuscript has been revised to add new updates to the ontology. The format has also been revised.

